# Design and development of a field-deployable water bath for loop-mediated isothermal amplification assay

**DOI:** 10.1101/2024.05.14.594127

**Authors:** Nafisa Rafiq, Mohit S. Verma

## Abstract

Nucleic acid testing has become a prominent method for rapid microbial detection. Unlike polymerase chain reaction (PCR), loop-mediated isothermal amplification (LAMP) is a simple method of nucleic acid amplification where the reaction can be performed at a constant temperature and the output provided in a colorimetric format. A transparent water bath is a desirable instrument to perform the heating and observe the visual results. However, existing methods of heating water are not convenient for loading and unloading the test samples. Here, we developed a field-deployable water bath—an isothermal heater called IsoHeat for short–which is solely dedicated to performing LAMP reactions and can heat the water up to 85°C. Using 3D-printing and laser-cutting technology, we fabricated different parts and mechanically assembled the parts to develop the entire device. Users can commence the heating by pressing the start button on the screen after entering the target temperature. Subsequently, the device heats up the water bath and maintains the target temperature through a PID algorithm-based control system. We demonstrate that IsoHeat can operate in environmental temperatures ranging from 5-33 °C and it can conduct LAMP reactions in liquid format as well as in paper-based devices. IsoHeat is more efficient and user-friendly compared to a commercially available immersion-heating device, which is often used to perform LAMP reactions. This newly developed device would be helpful to detect pathogens conveniently in the field (e.g., at point-of-care for human applications, on farms for plant and animal applications, and in production facilities for food safety applications).

## Introduction

Loop-mediated isothermal amplification (LAMP) (Notomi et al., 2000; Nzelu et al., 2019; Sirichaisinthop et al., 2011) is an isothermal molecular method for the rapid detection of nucleic acids (Conrad et al., 2020; Mohan et al., 2021). Although LAMP is not as common as polymerase chain reaction (PCR) for detecting pathogens, it can accomplish the detection in under an hour and does not require a thermal cycler (Kozel and Burnham-Marusich, 2017). In a LAMP assay, the samples are heated at a constant temperature for around 60 minutes, which causes the amplification of the nucleic acid (Mori et al., 2004; Nagamine et al., 2002; Tomita et al., 2008). Based on the presence of the target gene from the microbe, a change in fluorescence, turbidity, or color is observed after heating for less than an hour (Mori et al., 2001; Parida et al., 2008). To enable the monitoring of color change during LAMP with naked eyes (which could enable the determination of the quantity of the target), the sample should be heated in a transparent chamber. Moreover, while heating the nucleic acid sample, the temperature control should be precise, and the device should be convenient to use (Liu et al., 2017; Wang et al., 2011). There are many water-based heating devices available in the market, which can be used to heat the test samples. Circulating water baths have been used by researchers to maintain a constant and uniform temperature for biological assays (Tyler et al., 2009; Wang et al., 2006).

However, most of the commercially available water baths have non-transparent chambers to heat the water. In addition, although researchers have been using different types of precision heating devices to heat water while performing LAMP (Kellner et al., 2020; Pascual-Garrigos et al., 2021; Peltzer et al., 2021), they require the user to immerse their hands in hot water to place the samples.

Therefore, we have worked on developing a field-deployable transparent water bath that does not require the user to touch the hot water while placing the samples and allows tracking of the nucleic acid amplification visually. We aimed to make this system portable, having uniform heat distribution throughout the water while heating and allowing the user to observe continuous changes in color of the samples with naked eyes. Tracking the color changes over time has the potential to enable a quicker response for samples with higher concentrations of the target. It also has the potential to provide a quantitative response if appropriate image analysis algorithms are developed (outside the scope of the current work). Based on the sizing of the current water bath setups (Davidson et al., 2021; Pascual-Garrigos et al., 2021; Wang et al., 2021) used by researchers to perform LAMP, we aimed to keep our water container dimension within 13’’×13’’×8’’. Furthermore, we also intended to have a temperature error within ±0.5 °C since there are always some temperature fluctuations due to various factors and inherent challenges associated with thermal systems. Our transparent Isothermal heater (IsoHeat) can be used to perform LAMP assay in non-lab spaces and make it more user-friendly.

A step of LAMP is to heat the sample over a specific a period of time (typically less than one hour) at a constant temperature to facilitate nucleic acid amplification. The amplification of target nucleic acid can be observed visually as a change in turbidity or color (Mori et al., 2001; Parida et al., 2008). According to (Parida et al., 2008), this amplification can be done using a heating block and/or water bath. To our knowledge, there is no published work on the development of a water bath heating device dedicated to performing LAMP, which is convenient to use and allows visualization of amplification with the naked eye or a simple camera. Our group has previously used a commercially available precision cooker for heating the water to perform the LAMP assay (Pascual-Garrigos et al., 2021). That experimental setup was not sealed and required taping of the sample tubes to the wall of the water bath. However, taping the samples inside a hot water container is not convenient for the user, as it requires the use of heat-resistant gloves, which reduce the dexterity of operation. Arumugam *et al*., (Arumugam et al., 2020) also used the Anova immersion heater and transparent food storage container to create the water bath setup for COVID-19 detection using RT-PCR. Their water bath setup was exposed to air and they used a Raspberry Pi controlled servomotor to adjust and move the PCR tubes. Here, we developed a completely sealed water bath heating system where the user can hang the samples without immersing their hands in the hot water.

Table 1 shows a brief comparison of our developed heater IsoHeat with the existing heating devices. In comparison with the published water heating devices, the newly developed water bath has two distinguishing features: i) a sealed container for uniform heating and ii) a holder for hanging samples. These features make our device more convenient to use. IsoHeat is also cost-effective (Table S1) and allows heating the water in a short time (around 12 minutes).

**Table 1:** Comparison among different heating devices.

|  | **IsoHeat**<br>**(current work)** | **Anova Culinary**<br>**AN500-US00 Sous**<br>**Vide Precision**<br>**Cooker (ANOVA,**<br>**2023)** | **CORIO™ Open**<br>**Heating Bath**<br>**Circulator**<br>**1181J97**<br>**(Scientific, 2023)** | **Precision™**<br>**General**<br>**Purpose**<br>**Baths**<br>**TSGP02**<br>**(Fisher,**<br>**2023)** |
| --- | --- | --- | --- | --- |
| **Transparency**<br>**of water**<br>**chamber** | Transparent | Depends on the<br>water container<br>being used | Transparent | Not<br>transparent |
| **Heating**<br>**system** | Closed heating<sup>a</sup> | Open heating<sup>b</sup> | Open heating<sup>b</sup> | Closed<br>heating<sup>a</sup> |
| **Capacity** | Water capacity<br>is 6L | Variable capacity<br>depending on the<br>container size | Water capacity is<br>5L | Water<br>capacity is<br>2L |
| **Cost** | Development<br>cost of the water<br>bath is around<br>\$163, additional<br>\$47.5 required<br>for the Sample<br>holding<br>accessories | Heating device cost<br>\$150, additional<br>costing required for<br>container and extra<br>parts | Device costs<br>\$2272.08 | Device costs<br>\$1050 |

<sup>a</sup> Closed heating systems cannot exchange any matter (i.e., vapor)<sup>b</sup> Open heating systems can exchange matter

## Materials and Methods

Fig 1(a) shows the schematic of the device. The device consists of three major parts: the electrical control system, the hardware, and the power supply unit. The primary purpose of this device is to heat the water to a desired temperature and maintain it as long as required to perform the LAMP assay, which is accomplished through the control system. The hardware includes the water bath container, the lid of the container and some accessories to make the performance smooth and efficient. In our design, the lid holds the entire control system, sample loading unit and other accessories such as LCD display (B07KKB5YS9, Osoyoo), rotary motor (B08RWP6GJF, Cold and Colder), and sample holders. When the water bath is turned on and a target temperature is set on the display user interface, the heating element starts heating the water. With the help of the solid-state relay, the microcontroller maintains the desired temperature by turning the heating element on and off.

**Figure 1.**
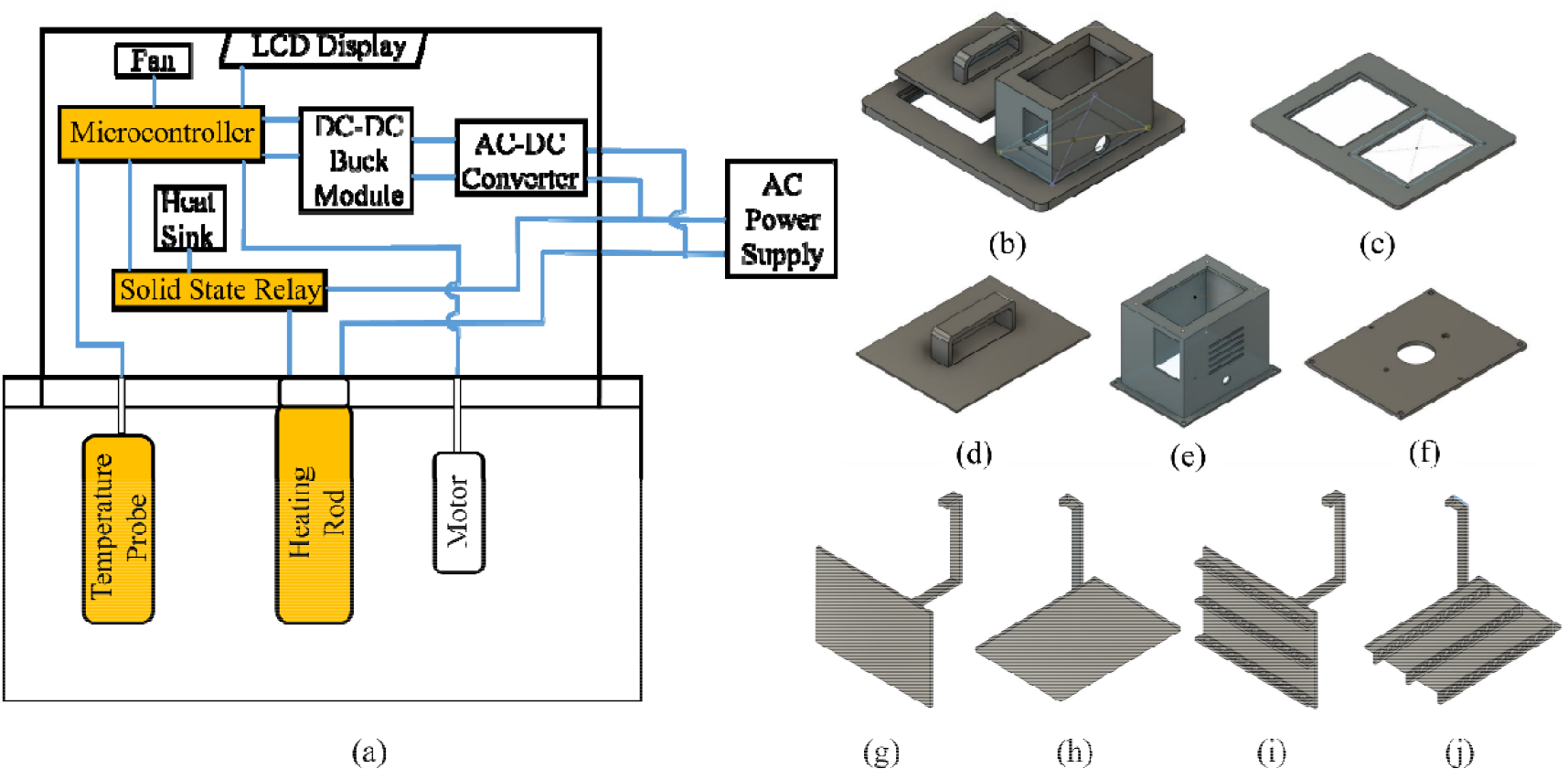
(a) Schematic model of the water bath. The 3D models of all parts of the lid, (b) The assembled lid, (c) the base of the lid, (d) the cover with loading slot, (e) the container for holding the electronic components, (f) the plate for holding the heating element. Accessories to hold the DNA sample. (g) Holder to carry paper samples from side, (h) holder to carry paper samples from, bottom (i) holder to carry both paper and/or liquid samples from side, (j) holder to carry both paper and/or liquid samples from bottom

### Electronic Control System

The control system performs the role of managing and controlling the behavior of other systems. It consists of a Raspberry (B07TD43PDZ, Raspberry Pi Foundation, Cambridge, England, UK), a solid-state relay (B01N0L5WSU, HiLetgo), a heating element of 1650W-120V (B07NPZ7LHX, Camco), and a temperature sensor (B09NVFJYPS, Bojack), designated by yellow-colored boxes in Fig 1. In our device, we used a closed-loop control system, which maintains the desired temperature of the water based on the feedback given by the temperature sensor. The Raspberry Pi microcontroller receives data from the temperature sensor containing the current temperature of the water and sends signals to the solid-state relay to allow or block the flow of current to the heating element based on the current temperature reading and the target temperature. Through this process, the system reaches a target temperature and maintains it for an indefinite amount of time.

### Hardware

In our heater, we used a transparent container of 12.8’’×10.4×6’’ (B00D0QSWX8, Carlisle) to hold the water. We preferred to use a commercially available container as developing a transparent container may require injection-molding technology, which might not be cost-effective to build a single prototype. The novel part of our device is the lid, which consists of the base, control system container, a plate to hold the heating rod and a cover with a sample hanging slot. Fig 1(b-f) illustrates the three-dimensional (3D) models of the parts made with Autodesk Fusion 360. The base of the lid was made with plywood coated with a waterproof solution. We used Trotec Speedy 400 Laser Cutter to cut out the base from a 24’’× 24’’×0.25’’ plywood board (5420278, Ace Hardware) (Fig S1). The control system container was made with Raise 3D printer using acrylonitrile butadiene styrene (ABS) filament (B08CC7QFXM, NovaMaker) (Fig S3). The plate and the sample loading cover were fabricated with Formlabs Form 3+ Stereolithography (SLA) 3D printer using High Temp Resin V2 (RS-F2-HTAM-02, Formlabs) (Fig S2 and S4).

To insert the tubes containing DNA samples inside the water bath, we fabricated multiple sample holders with Rigid 4000 V1 Resin (RS-F2-RGWH-01RS, Formlabs) using the same SLA 3D printer, which is suitable for paper-based and liquid-based LAMP reactions (Fig S5-S8). We used orf7ab.I primer mix (Davidson et al., 2021) to prepare the master mix for detecting the orf7ab segment of SARS-CoV-2 genome. We used a synthetic DNA target for this segment for ease of handling. Table S2 describes the sequences for the primers. The concentration for the orf7ab DNA was 10^6^ copies per reaction for the liquid colorimetric LAMP and 5×10^6^ copies per reaction for the paper-based colorimetric LAMP. The sample holders for carrying paper LAMP samples from the side and bottom of the water bath are shown in Fig 1(g and h), respectively, whereas Fig 1(i and j) shows the sample holders for holding small tubes which can carry both paper and liquid LAMP samples from the side and bottom, respectively. We used a submergible rotary motor to circulate the water and maintain a uniform temperature all over the water bath.

Moreover, a touchscreen LCD display connected to the microcontroller displays the current temperature of the water and allows the user to set a target temperature and start/stop the heating process (Fig 2a). The final images of the device are shown in Fig 2(a-d), where the other parts are assembled properly, except the sample holders.

**Figure 2.**
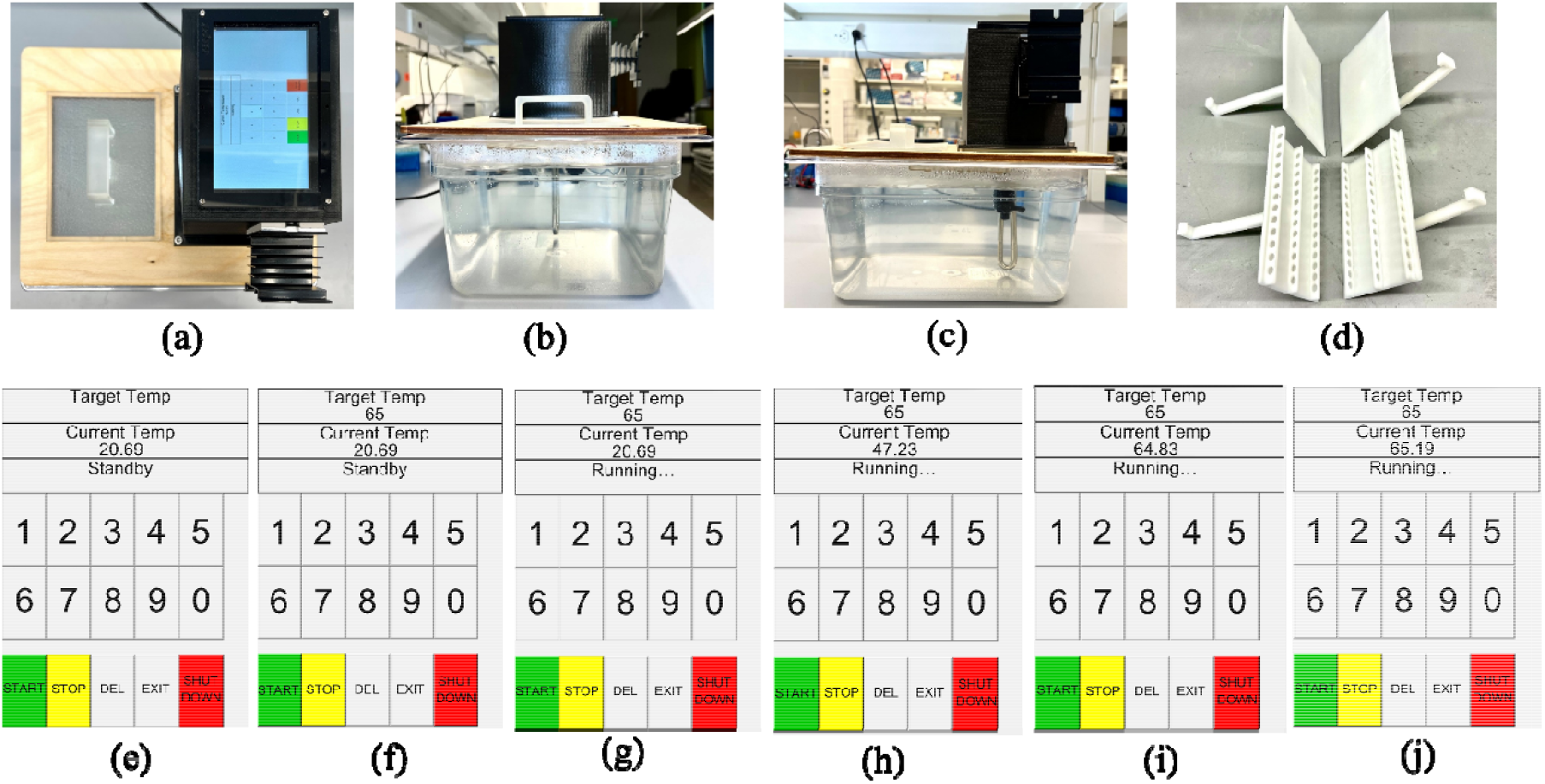
Images of water bath heater with sample holding accessories, (a) Top view, (b) front view, (c) side view of the water bath and (d) sample holders to carry the samples. Screenshots of software operations-(e) in standby mode, (f) in standby mode after setting the target temperature, (g) after pressing “start”, (h) halfway through reaching the target temperature, (i and j) maintaining the target temperature.

### Power supply unit

We designed a power supply unit to run the electronic components and the heating element from a single power source. The heating element was directly connected to a 120V AC power outlet, whereas an AC-DC converter (B0BB8YWBHX, Diann) and a DC-DC buck module (B01HXU1NQY, HiLetgo) were connected in series to convert the 120V AC supply to 6V DC and supply power to the Raspberry Pi. All other electronic components were powered by the Raspberry Pi.

## Results and Discussion

### Temperature fluctuations from the target temperature

The typical heating temperature for LAMP is 65 °C (Davidson et al., 2021; Mohan et al., 2021; Parida et al., 2008; Pascual-Garrigos et al., 2021; Safavieh et al., 2014; Velders et al., 2018; Wan et al., 2019; Wang et al., 2023, 2022, 2021; Zhang et al., 2019). Therefore, we tested the heating efficiency of the water bath by setting 65 °C as the target temperature. We tested the heater in an environment set to room temperature (21 °C) and observed the heating. We also tested the water bath in different environments, such as inside a refrigerator (5 °C) and outdoors on a hot day (33°C). Fig 3(a) shows the temperature curves with time in three different environments.

**Figure 3.**
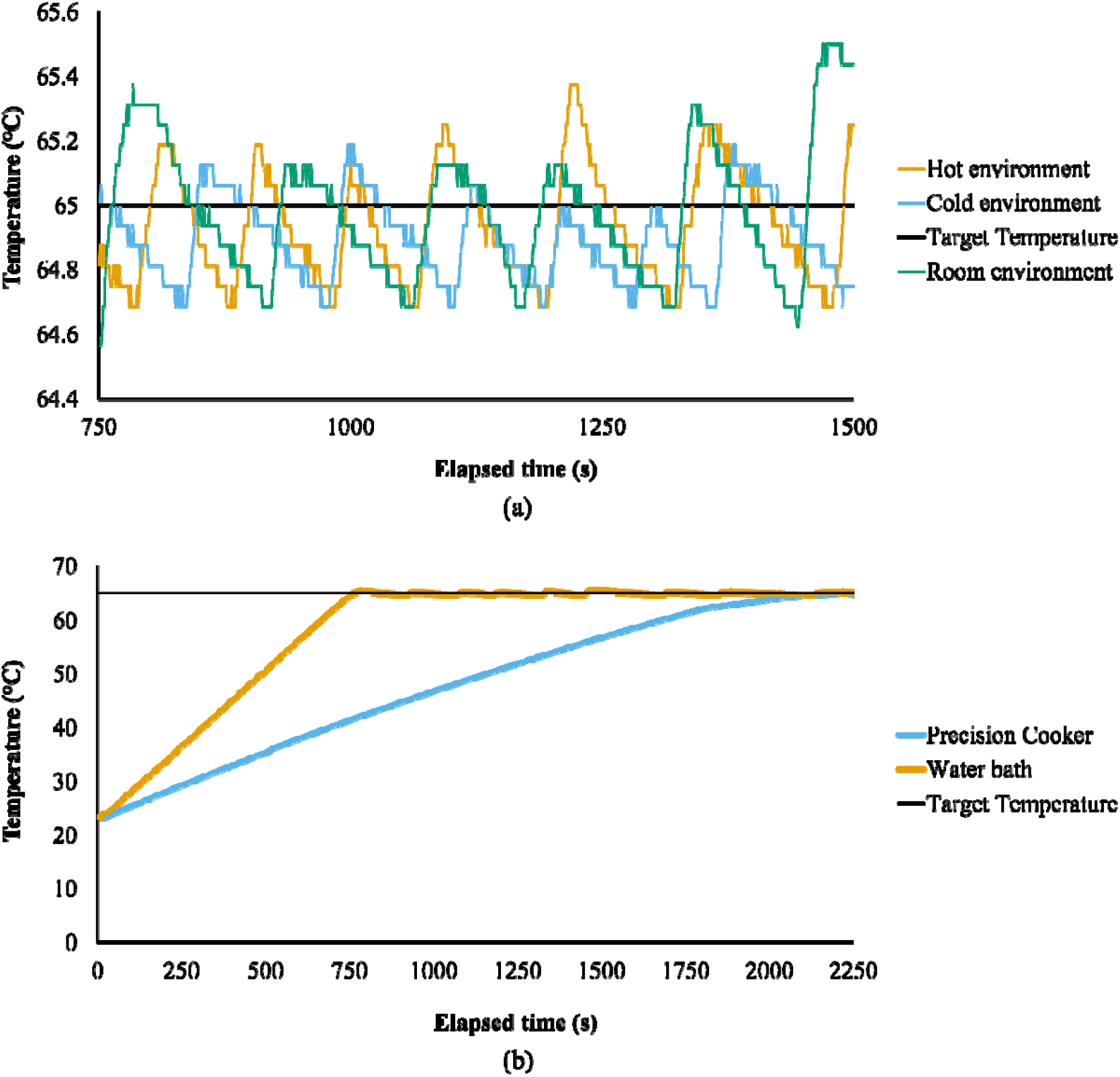
(a) Variation in temperature fluctuation in different environments after reaching the target temperature, (b) Variation of time for precision cooker and water bath to reach the target temperature while heating 6L of water.

Considering the temperature curve from the aforementioned three environments, Fig 3(a) illustrates that the temperature fluctuation in these environments was no greater than ± 0.5°C and IsoHeat performs similarly in all three environments.

### Validation of temperature through IR images

After reaching the target temperature initially, the control system maintained the temperature by turning the heating element on and off. We examined the validity of the projected temperature using an Hti HT-04 Thermal Imaging Camera. As mentioned in the previous section, when the temperature was set to 65°C, the fluctuation was within ±0.5°C. Infrared images of different points of the water bath were taken with the thermal imager. Fig 4(a) illustrates the thermal images of various points when the target temperature was set to 65°C. The minimum and maximum observed temperatures through the thermal imager were 64.4°C and 65.1°C, respectively. Fig 4(b) shows the IR images of a single point over time, where the lowest detected temperature was 64.8°C and the highest temperature was 65.8°C. Since the thermal imager has an error margin of ±1°C and a thermal sensitivity of 0.07°C, the thermal images confirmed the temperatures recorded by the temperature sensor in the water bath.

**Figure 4.**
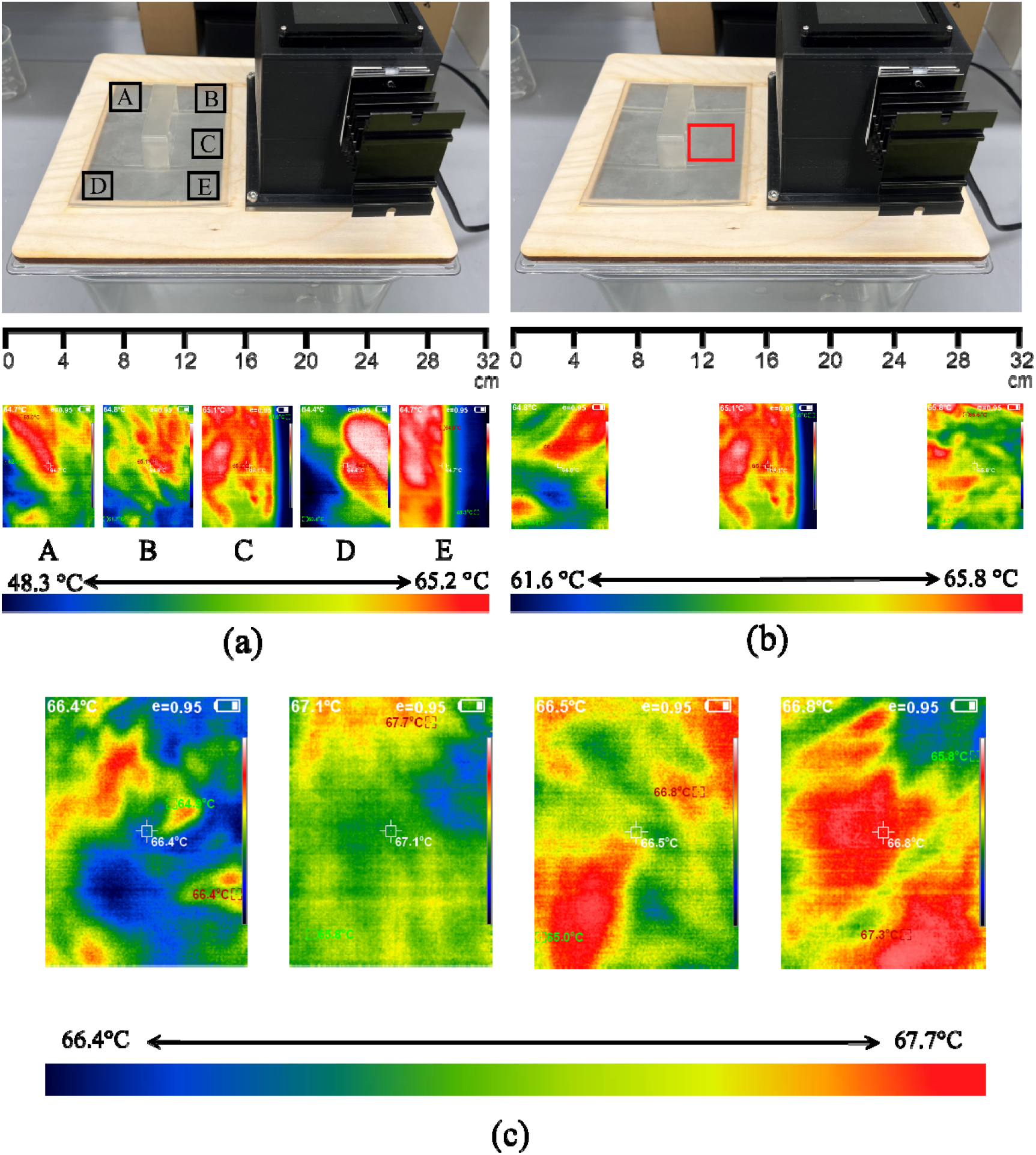
Thermal images of (a) different points of water after reaching the target temperature (65°C), (b) a single point at arbitrary time after reaching the target temperature (65°C), (c) the water heated with the Anova precision cooker after reaching the target temperature (65°C)

### Visual observation of LAMP assays

The primary objective of our work was to develop a water bath heater that can be used to heat the colorimetric LAMP samples and illustrate visible results. We performed both the paper-based and liquid-based colorimetric LAMP assays using the water bath and received satisfactory results. In paper based colorimetric LAMP, the samples are added to a paper-device (Davidson et al., 2021), whereas an individual domed PCR tube holds the samples in liquid-based colorimetric LAMP (Wang et al., 2023). We used a DNA target for simpler handling as compared to RNA that is naturally found in SARS-CoV-2. Before placing the sample inside the water bath, we turned on the device by connecting it to the power port and set the target temperature to 65°C. It takes around 12 minutes to reach 65°C when the water chamber is filled with water of its capacity, 6L. Once the water reached the target temperature, the samples were loaded in the 3D-printed sample holders and placed inside the water bath. Fig 5(a and b) and Fig 5(c and d) illustrate the heating and color change of the samples for liquid-based and paper-based colorimetric LAMP, respectively, where we used the sample holder compatible for the liquid-based LAMP as well as paper-based LAMP. Fig 5(e and f) shows the heating of paper-based LAMP, where the samples were placed inside Ziploc bags and taped to the sample holders. This experimental setup is compatible with heating larger sizes of paper-based LAMP. In all cases (Fig 5), visible color change was observed in the positive control samples after heating for 60 minutes, which indicated the presence of the specific nucleic acid. On the contrary, no colorimetric change occurred in the negative control samples even after the incubation.

**Figure 5.**
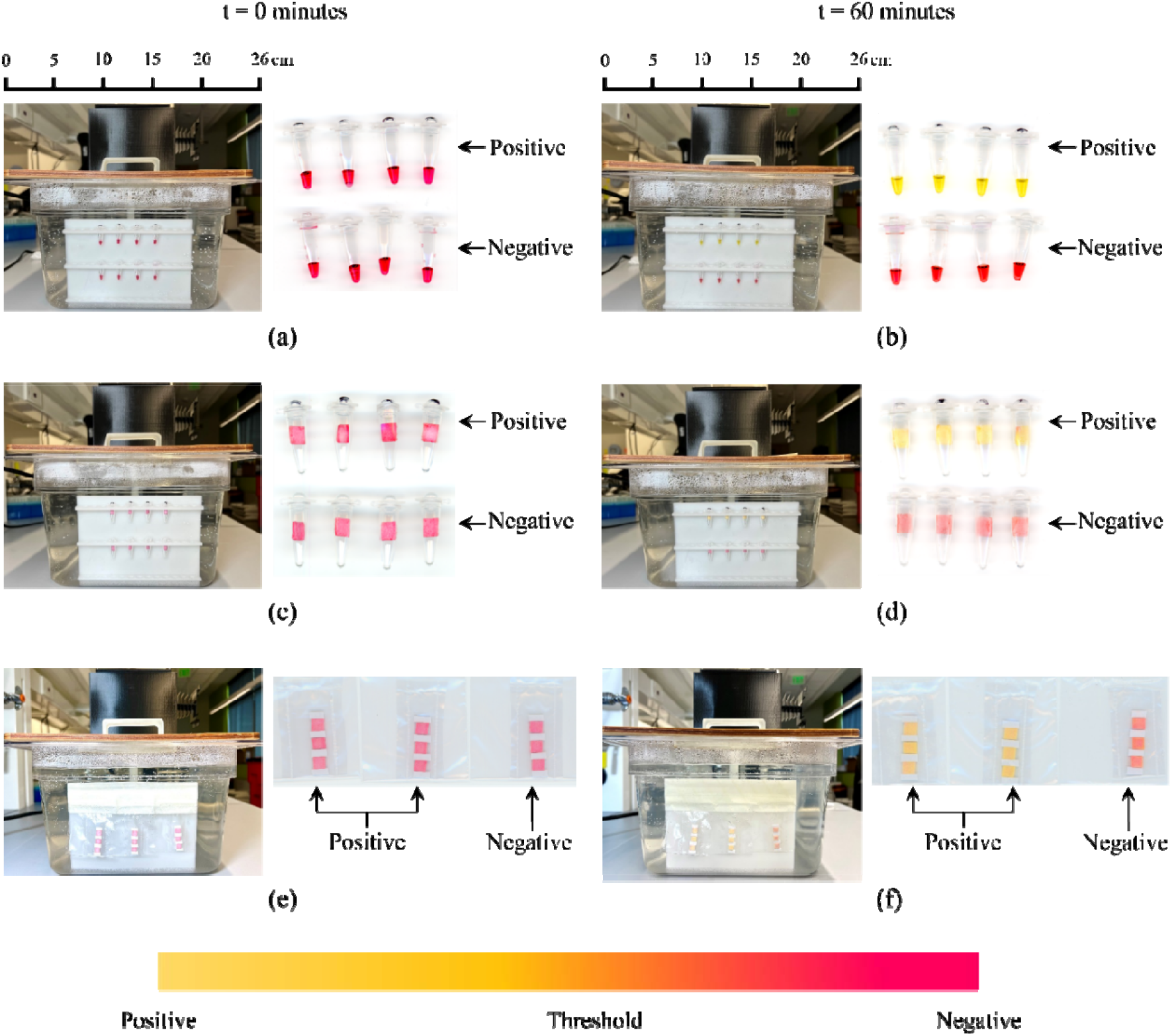
Performing colorimetric LAMP using the water bath. Positive samples have the presence of the target DNA and negative samples have nuclease free water instead of the target DNA. Tube holders holding the liquid samples inside the water bath at (a) t=0 minute with scanned image of the samples before heating and (b) t=60 minutes with scanned image of the samples after heating for 60 minutes. Tube holders, holding the paper samples inside the water bath at (c) t=0 minute with scanned image of the samples before heating and (d) t=60 minutes with scanned image of the samples after heating for 60 minutes. Sample holders holding the paper samples inside the water bath at (e) t=0 minute with scanned image of the samples before heating and (f) t=60 minutes with scanned image of the samples after heating for 60 minutes

### Comparison with Anova precision cooker

Based on the literature, one of the most common heating equipment used for heating LAMP assays in the field is *sous vide* precision cooker or *sous vide* immersion heater (Davidson et al.,

2021; Kellner et al., 2020; Pascual-Garrigos et al., 2021; Peltzer et al., 2021; Wang et al., 2023, 2021). We tested our device, the “IsoHeat” and the Anova *sous vide* precision cooker in identical environments to make a brief comparison between them. Fig 3(b) exhibits the temperature curve over time when 6L water was heated using the two devices. Our developed water bath required around 12 minutes to reach the target temperature (65 °C) for the first time, whereas the precision cooker took around 36 minutes to reach the same temperature. Moreover, the accuracy of water temperature heated with the precision cooker was also questionable. Fig 4(c) presents the IR images of the water heated with the precision cooker after reaching the target temperature of 65°C. On the other hand, temperature fluctuation in the IsoHeat is shown in Fig 4a, when the target temperature was set to 65°C. Even though the IsoHeat is more time efficient and has higher accuracy than the precision cooker, the manufacturing cost of the lid is comparable to the price of the Anova precision cooker. The Anova precision cooker costs $150 and an additional cost of $12 is required to create the heating setup using the commercially available transparent container. On the other hand, IsoHeat costs around $160 to fabricate including the water container, which is similar to the cost of the aforementioned setup.

## Conclusion

Here, we developed a water bath system that is capable of heating the LAMP assays in any confined or open place and is able to generate results visible to the naked eye. Here, we have achieved the goals that we aimed for, namely: i) the system is portable (water container dimensions are 12.8’’×10.4 × 6’’), making it suitable for on-farm diagnostics, ii) it maintains uniform heat throughout the system (the temperature fluctuation is within ±0.5 °C) and iii) continuous color change can be observed with naked eyes as it is transparent. Other advantages of our device are the sample loading and unloading do not require direct touch of the hot water with the users’ skin, therefore, safe to use and the estimated development cost of the entire setup is ~$210.

Although the developed water bath features fast heating and minimal temperature fluctuation, it has some limitations. Firstly, when the water bath is filled with water, it becomes a little heavy, which may adversely affect its portability. However, without any water, the device is lightweight and easily portable. Secondly, the water bath is unable to maintain the exact target temperature, with fluctuations of ± 0.5 °C. However, from our experience, LAMP reactions did not show any sensitivity to ± 0.5 °C temperature fluctuation.

Future work can focus on improving the functionality of the device by reducing temperature fluctuation and increasing accuracy. A compact and lightweight water bath heater can be developed to perform the heating of LAMP assays, which will increase the portability of the device. Moreover, instead of using a commercially available water container, a custom-made container can improve the aesthetics and make it more user-friendly. Innovations in materials and design can lead to more robust, corrosion-resistant, and user-friendly heating devices. This could also entail better cleaning and maintenance procedures. To maintain user safety and adherence to industry rules, it could be a priority to continuously develop safety features including overheat protection, leak detection, and automatic shutdown methods.

## Acknowledgments

We are thankful to our lab members Jiangshan Wang for reviewing the manuscript, Gopal Palla for providing colorimetric LAMP training, Josiah Levi Davidson for providing the LAMP primers and Mohsen Ranjbaran for explaining the importance of heating elements in colorimetric LAMP.

## Declaration of AI and AI-assisted technologies in the writing process’

During the preparation of this work the authors used Grammarly (https://grammarly.com/) and ChatGPT (https://chat.openai.com/) in order to check for grammar errors and improve the academic writing language. After using this tool/service, the authors reviewed and edited the content as needed and take full responsibility for the content of the publication.

