## Supporting Information for "Design and development of a field-deployable water bath for loop-mediated isothermal amplification assay"

**For**

1.1. *Supplementary Figures*

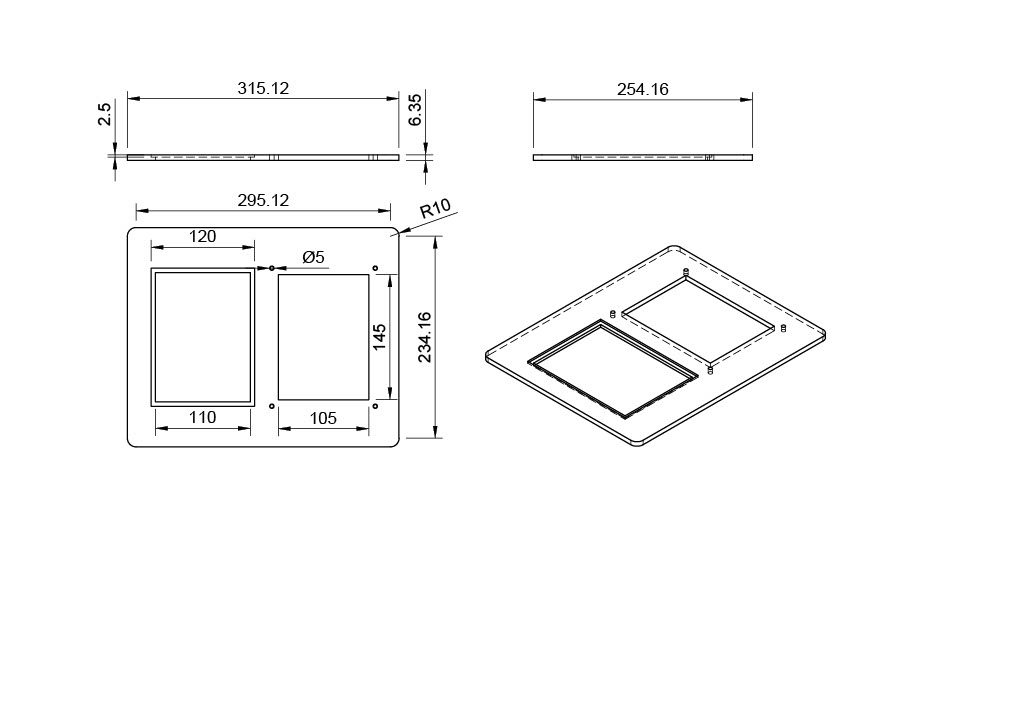

Figure S1. Design details of the base of the lid, shown in Figure 2 (b)

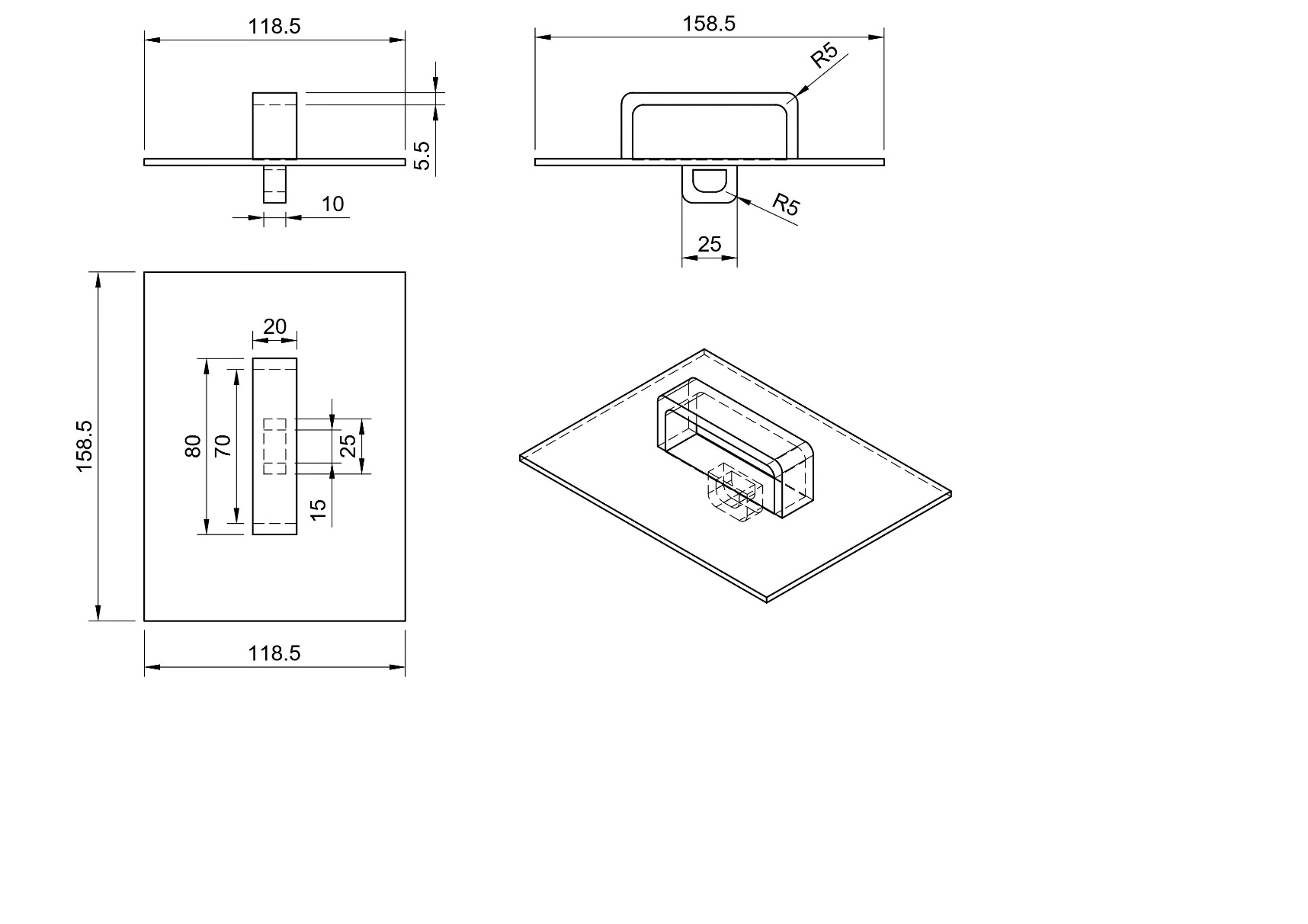

Figure S2. Design details of the sample loading lid cover, shown in Figure 2(c)

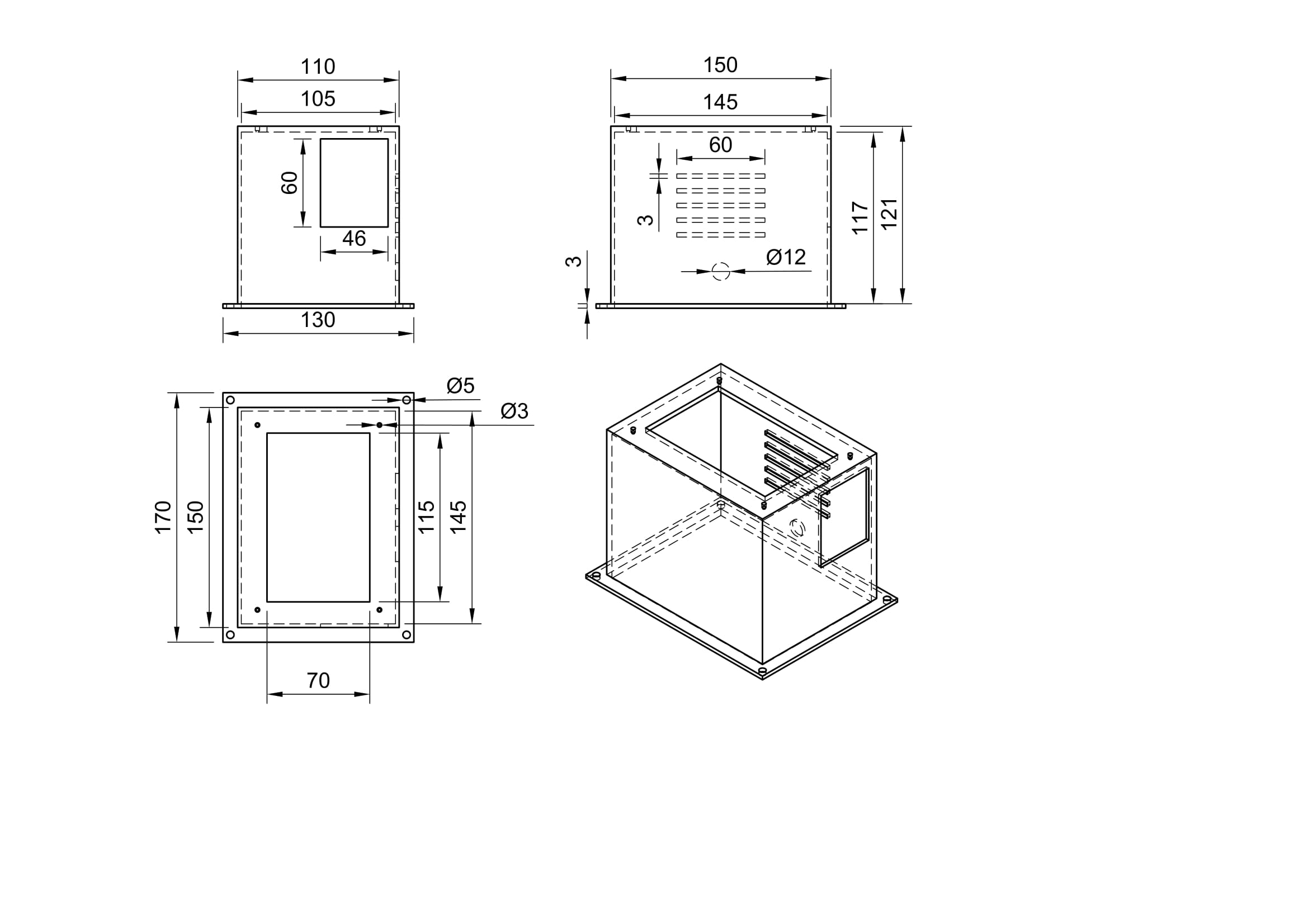

Figure S3. Design details of the container for holding the electronic components, shown in Figure 2(d)

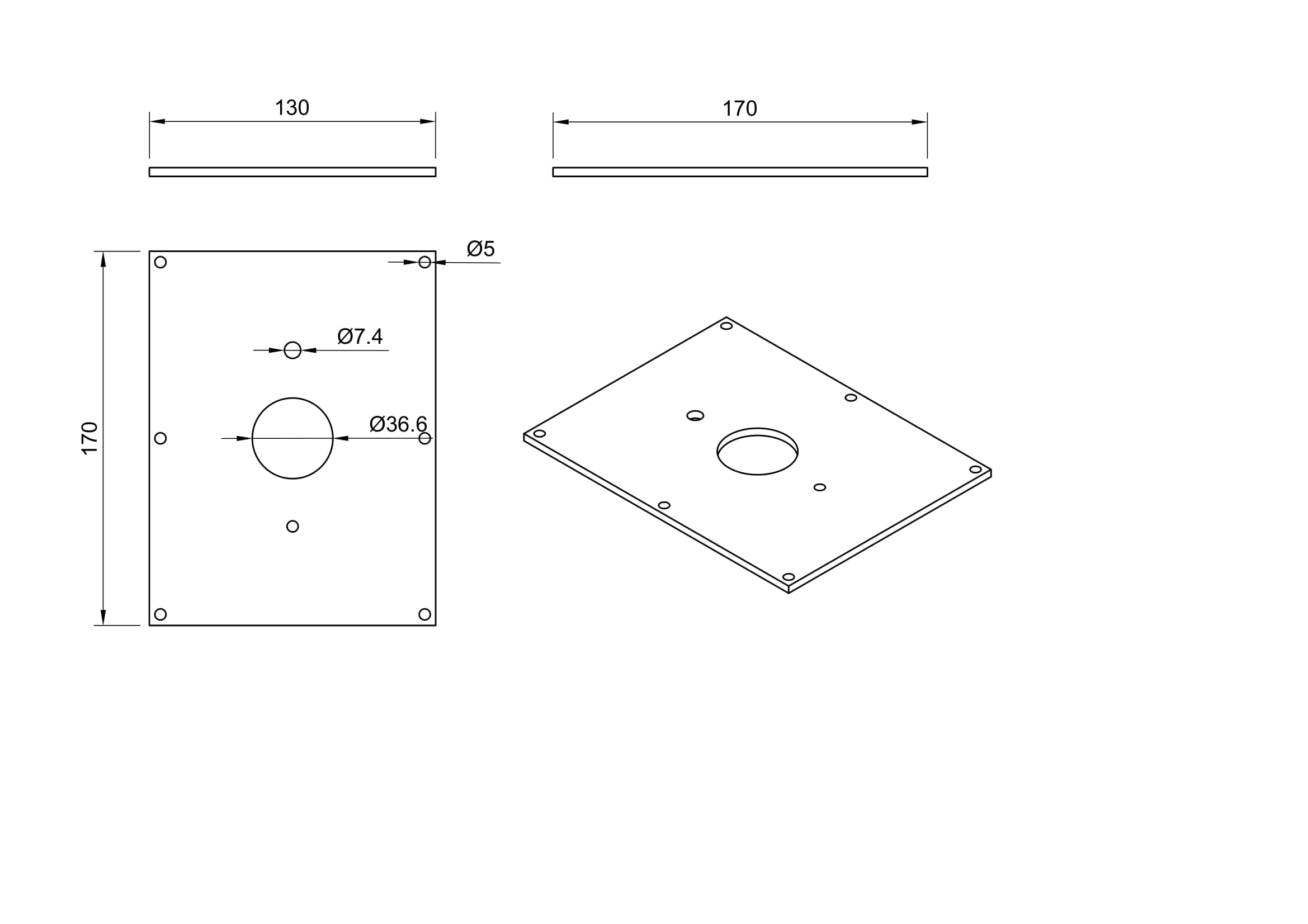

Figure S4. Design details of the plate for holding the heating element, shown in Figure 2(e)

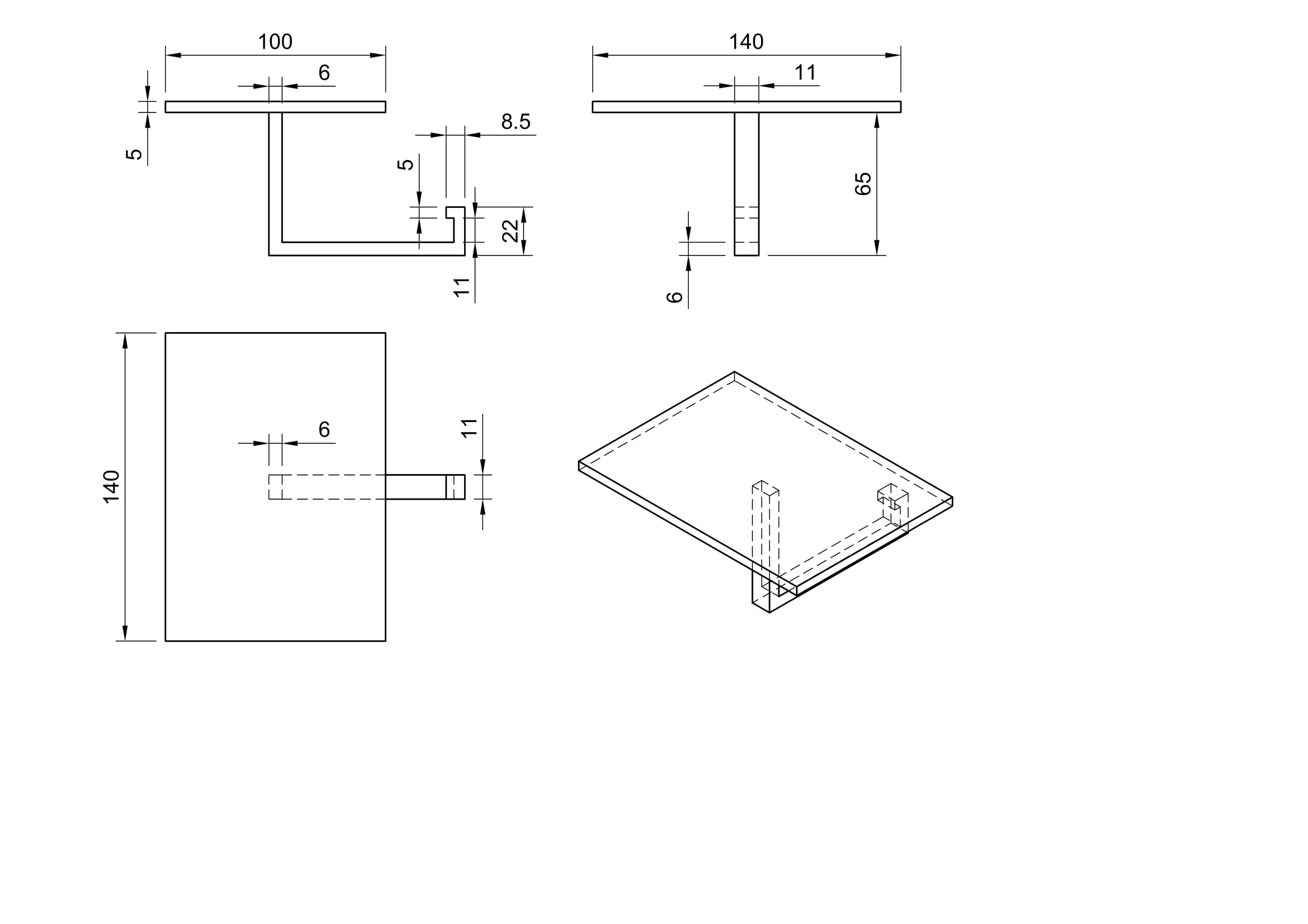

Figure S5. Design details of the holder to carry paper samples from side, shown in Figure 3(a)

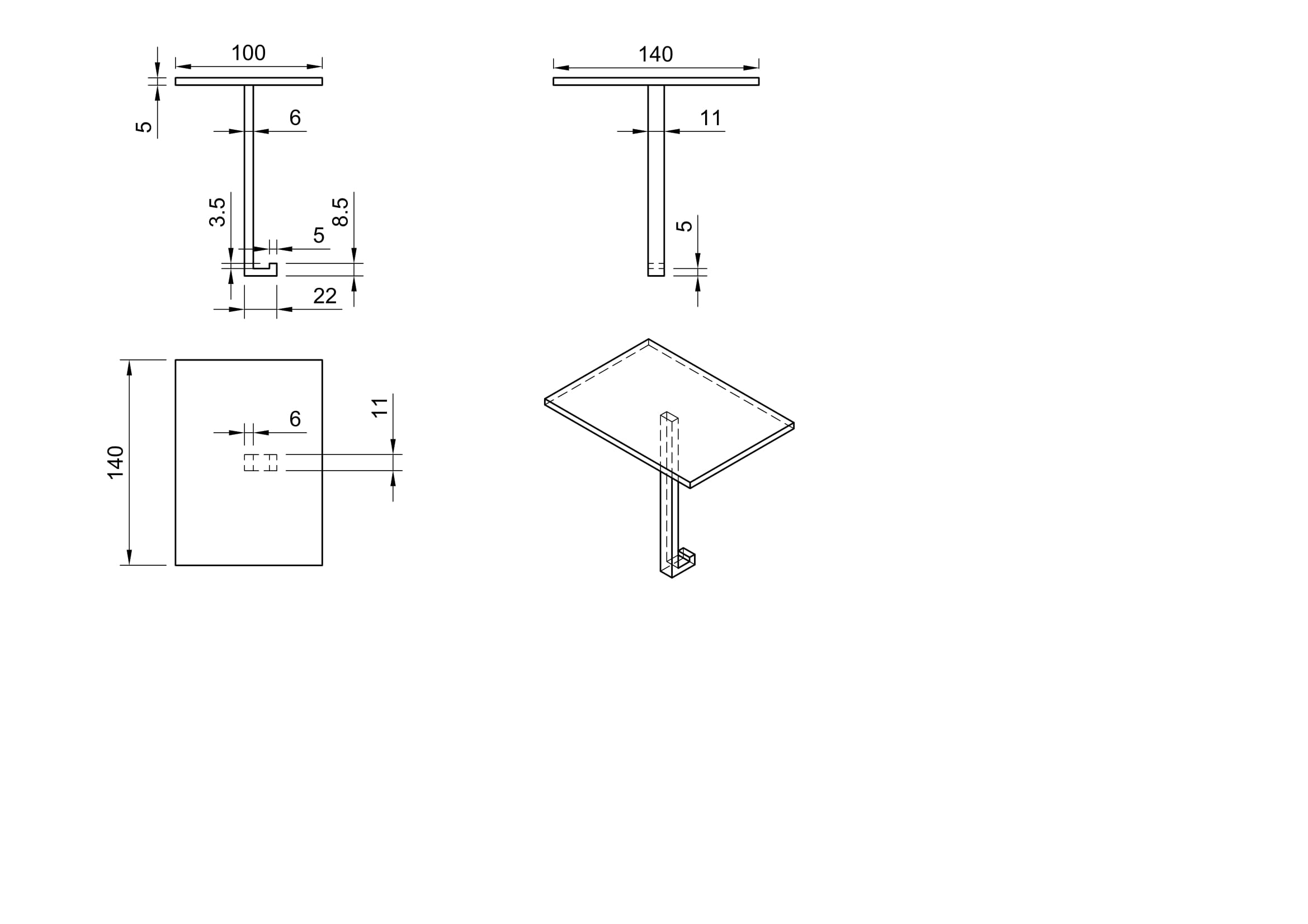

Figure S6. Design details of the holder to carry paper samples from bottom, shown in Figure 3(b)

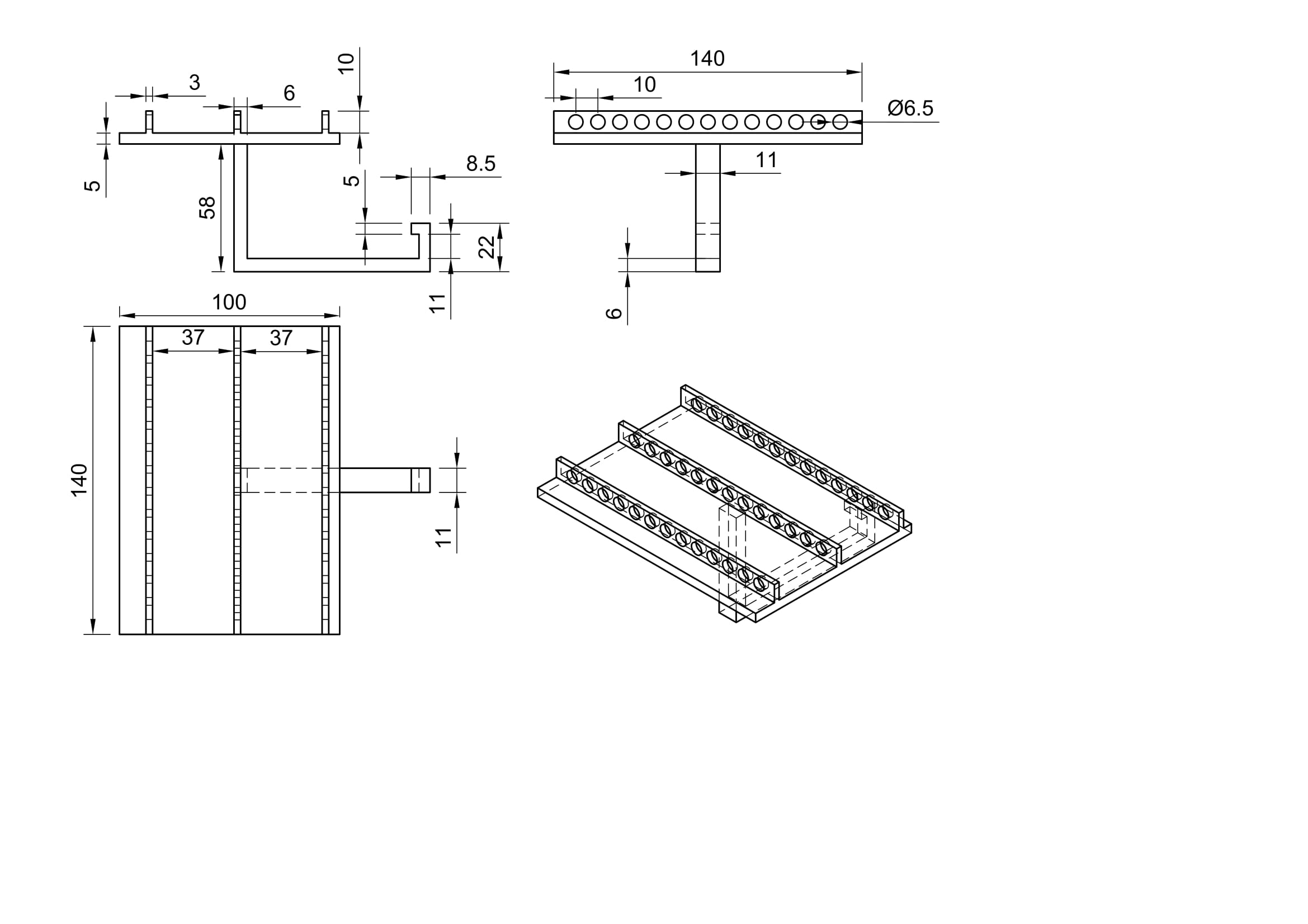

Figure S7. Design details of the holder to carry samples from side, shown in Figure 3(c)

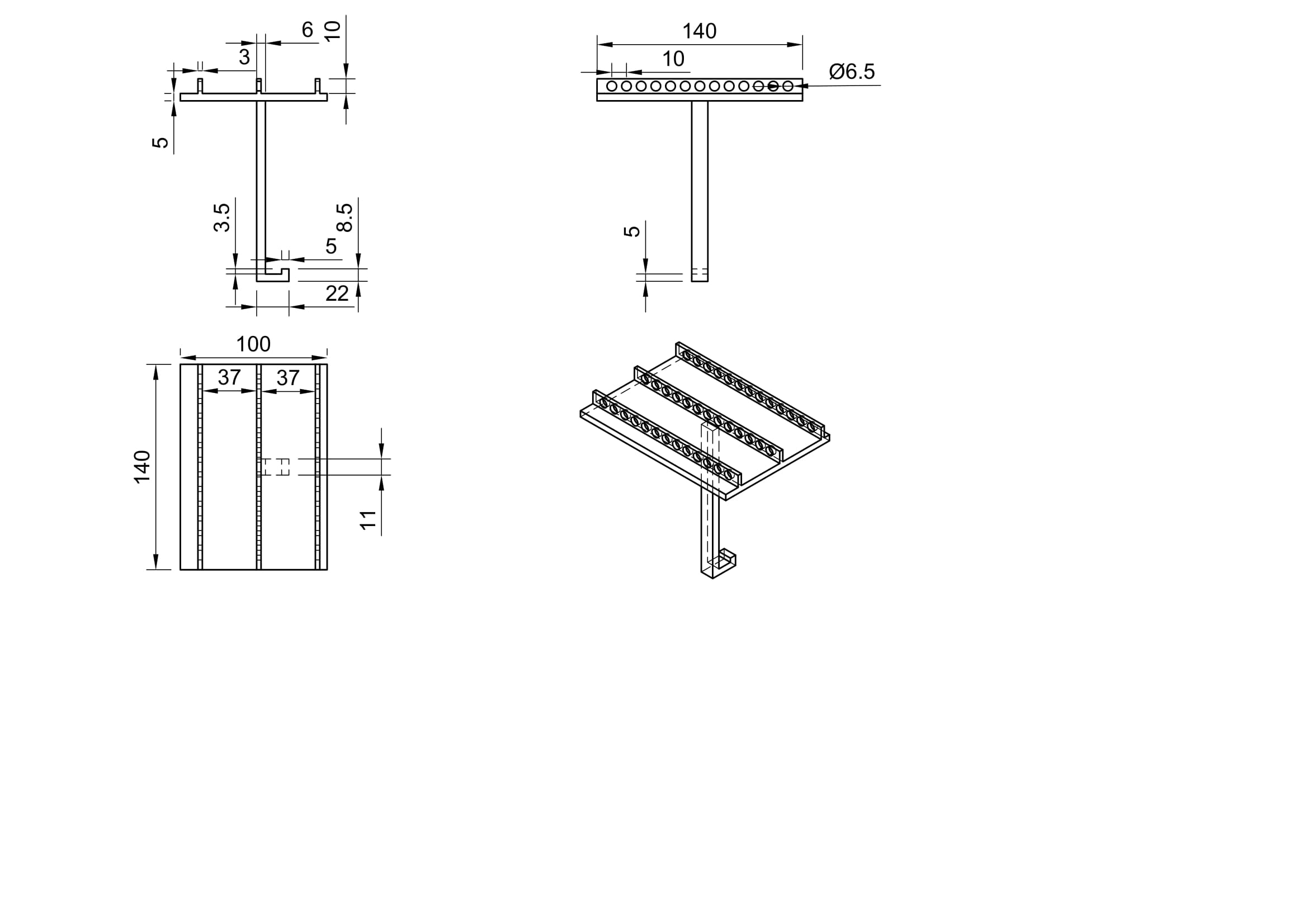

Figure S8. Design details of the sample holder to carry samples from bottom, shown in Figure 3(d)

1.2. *Supplementary Tables*

Table S1: Bill of materials used to develop the heating system (Prices are based on academic laboratory prices from vendors)

| **Item** | **Supplier** | **Catalog number** | **Quantity** | **Unit** | **Price ($)** | **Number of units used/ device** | **Price ($)/unit used** | **% Contribution to total cost** |
| --- | --- | --- | --- | --- | --- | --- | --- | --- |
| **Heating Device** | | | | | | | | |
| Raspberry pi microcontroller | Raspberry Pi Foundation, Cambridge, England, UK | B07TD43PDZ | 1 | Piece | 70 | 1 | 70.00 | 33.21 |
| SSR-40DA | HiLetgo | B01N0L5WSU | 2 | Piece | 10.5 | 1 | 5.25 | 2.49 |
| 1500W-120V Heating element | Camco | B07NPZ7LHX | 2 | Piece | 8.99 | 1 | 4.5 | 2.13 |
| DS18B20 Temperature sensor | Bojack | B09NVFJYPS | 2 | Piece | 5 | 1 | 2.50 | 1.19 |
| Silicon wires | BNTECHGO | B01MXGNTA4 | 50 | feet | 6.98 | 3 | 0.42 | 0.20 |
| Jumper wires | Edgelec | B07GD2BWPY | 120 | Piece | 6.99 | 8 | 0.46 | 0.22 |
| Aluminum Heat sink | Tihood | B07QP856MJ | 3 | Piece | 9.99 | 1 | 3.33 | 1.58 |
| Mini brushless water pump | Cold and colder | B08RWP6GJF | 1 | Piece | 11.6 | 1 | 11.6 | 5.50 |
| AC-DC buck converter | Diann | B0BB8YWBHX | 2 | Piece | 5.98 | 1 | 2.99 | 1.42 |
| DC-DC buck module | HiLetgo | B01HXU1NQY | 10 | Piece | 5.49 | 1 | 0.54 | 0.26 |
| Raspberry pi cooling fan | EasyCargo | ‎B0792BW2VH | 4 | Piece | 8.98 | 1 | 2.24 | 1.06 |
| ABS 3D Printer Filament, 1.75mm, Black | NovaMaker | B08CC7QFXM | 1 | kg | 14.99 | around 0.1 | 1.49 | 0.71 |
| Formlabs High Temp Resin V2 | Formlabs | RS-F2-HTAM-02 | 1 | L | $160 | Around 0.08 | 12.8 | 6.07 |
| Plywood board | Ace Hardware | 5420278 | 8 | Piece | 8.00 | 1 | 8.00 | 3.80 |
| 5’ LCD touch screen display | Osoyoo | B07KKB5YS9 | 1 | Piece | 28 | 1 | 27.16 | 12.89 |
| Carlisle 10222B07 storplus half size food pan | Carlisle | B00D0QSWX8 | 1 | Piece | 10.00 | 1 | 10.00 | 4.74 |
| **Sub-Total:** |  |  |  |  |  |  | **163.28** | 77.46 |
| **Sample Holders** | | | | | | | | |
| Formlabs Rigid 4000 Resin V1 | Formlabs | RS-F2-RGWH-01RS | 1 | L | $190 | Around 0.25 L | 47.5 | 22.54 |
| **Total:** |  |  |  |  |  |  | **210.78** |  |

Table S2: Sequences for LAMP primers used in this publication.

| **Primer** | **Sequence (5’ - 3’)** |
| --- | --- |
| orf7ab.I_F3 | CGGCGTAAAACACGTCTA |
| orf7ab.I_B3 | GCTAAAAAGCACAAATAGAAGTC |
| orf7ab.I_FIP | GGAGAGTAAAGTTCTTGAACTTCCTAGTTACGTGCCAGATCAG |
| orf7ab.I_BIP | TGCGGCAATAGTGTTTATAACACTATGAAAGTTCAATCATTCTGTCT |
| orf7ab.I_LF | TGTCTGATGAACAGTTTAGGTGAAA |
| orf7ab.I_LB | TTGCTTCACACTCAAAAGAA |

**Software Code:**

```
# heater_final.py

# Author = Nafisa Rafiq
# Maintainer = Nafisa Rafiq
# Email =
# Status = Completed
# Laboratory = Verma Lab
# Date = 08/10/2023

# Import Guizero for the GUI
from guizero import *

# Import pandas and datetime
# for logging and saving data
import pandas as pd
from datetime import datetime

# Import other required python packages
import sys
import os
import glob
import time
import RPi.GPIO as GPIO
import PID

###### GUI Commands (Start) ######

# For updating target temperature display
# after pressing a number button
def screenCapture(obj):
    # Global variables for this function
    global screen_capture, TargetTempDisplay

    # When a number is pressed, display this number
    # in the target temperature display box
    obj = str(obj)
    screen_capture += obj
    TargetTempDisplay.value = screen_capture

# For starting the system
def StartSystem():
    # Global variables for this function
    global StartSeconds, Temp_thres, now, filename
    global curr_time, data, df, StatusDisplay, TargetTemp
    global TargetTempDisplay, SystemOn, pid

    # Check if target temperature has been set or not
    if not TargetTempDisplay.value == "":
        TargetTemp = float(TargetTempDisplay.value)

        # Update status display
        StatusDisplay.value = ""
        StatusDisplay.value = "Running..."

        # Set PID parameters and
        # temperature tolerance threshold
        Temp_thres = TargetTemp - 0.5
        pid.SetPoint = TargetTemp
        pid.setSampleTime(0.01)

        # System start time
        StartSeconds = time.time()

        # Start logging data
        now = datetime.now()
        filename = str(now)
        filename = filename.replace(" ", "_")
        filename = filename.replace(":", ".")
        curr_time = str(now)
        curr_time = curr_time.replace(" ", "_")
        curr_time = curr_time.replace(":", ".")
        data = [[curr_time, 0, 0, 0]]
        df = pd.DataFrame(data, columns = ['DateTime', 'Time (s)', 'temp', 'TargetTemp'])

        # Start the system
        SystemOn = True

    else:
        # Instruct the user to
        # enter target temperature

        StatusDisplay.value = ""
        StatusDisplay.value = "Enter target temp!"
        SystemOn = False

# For stopping the system
def StopSystem():
    # Global variables for this function
    global TargetTemp, screen_capture, TargetTempDisplay, SystemOn

    # Reset global variables
    TargetTemp = 0
    screen_capture = ""
    TargetTempDisplay.value = screen_capture
    SystemOn = False

    # Turn OFF the relay and the heater
    GPIO.output(18, GPIO.LOW)

    # Update status display
    StatusDisplay.value = ""
    StatusDisplay.value = "Standby"

# For removing character from
# target temperature display box
def backSpace():
    # Global variables for this function
    global screen_capture, TargetTempDisplay

    try:
        # Remove the last character
        screen_capture = screen_capture[0:len(screen_capture)-1]
        TargetTempDisplay.value = screen_capture
    except:
        # Pass if no characters remaining
        pass

# For exiting the app
def exitApp():

    # Turn OFF the relay and the heater

    GPIO.output(18,GPIO.LOW)

    # exit the app
    sys.exit()

# For shutting down the
# raspberry pi
def Shutdown():

    # Turn OFF the relay and the heater
    GPIO.output(18,GPIO.LOW)

    time.sleep(1)

    # Shut down
    os.system("sudo shutdown -h now")

###### GUI Commands (end) ######

##### Functions for reading temperature (Start) #####

# For reading the raw file
# containing temperature information
def read_temp_raw():

    f = open(device_file, 'r')
    lines = f.readlines()
    f.close()
    return lines

# For converting raw temperature
# data to degree celcius
def read_temp():

    lines = read_temp_raw()
    while lines[0].strip()[-3:] != 'YES':
        # Check until data has been found
        time.sleep(0.2)
        lines = read_temp_raw()

    # Check for the string containing
    # the temperature
    equals_pos = lines[1].find('t=')

    if equals_pos != -1:
        # Return the temperature
        # in degree celcius
        temp_string = lines[1][equals_pos+2:]
        temp_c = float(temp_string) / 1000.0
        return temp_c

##### Functions for reading temperature (End) #####

##### Main Function (Start) (Run indefinitely) #####

def mainRun():
    # Global variables for this function
    global StartSeconds, CurrSeconds, now, curr_time, df
    global PID_thres, pid
    global TargetTemp, CurrTempDisplay
    global SystemOn, FirstStepPassed
    global app

    # Check if system is running
    if SystemOn:

        # Read temperature
        i = read_temp()

        # Get PID reading
        pid.update(i)
        output = pid.output

        # Log and save data to .csv file
        now = datetime.now()
        curr_time = str(now)
        curr_time = curr_time.replace(" ", "_")
        curr_time = curr_time.replace(":", ".")
        CurrSeconds = time.time() - StartSeconds
        data2 = [[curr_time, CurrSeconds, i, TargetTemp]]
        df2 = pd.DataFrame(data2, columns = ['DateTime', 'Time (s)', 'temp', 'TargetTemp'])
        df = df.append(df2, ignore_index=True)
        df.to_csv('data/' + filename + '.csv')

        # Check if target temp has been
        # reached for the first time or not
        if FirstStepPassed == False:
            # Do not use PID threshold,
            # use temperature threshold

            # Compare current temperature
            # to temperature threshold
            if i > Temp_thres:

                # Turn OFF the relay and the heater
                GPIO.output(18,GPIO.LOW)

                # Print current temperature and
                # heater status
                print(i, "Degree celcius, ", "Heater OFF, ", "PID: ", output)

                # Update current temperature display
                CurrTempDisplay.value = str(round(i,2))
                app.update()

            else:
                # Turn ON the relay and the heater
                GPIO.output(18,GPIO.HIGH)

                # Print current temperature and
                # heater status
                print(i, "Degree celcius, ", "Heater OFF, ", "PID: ", output)

                # Update current temperature display
                CurrTempDisplay.value = str(round(i,2))
                app.update()

            # Target temp has been
            # reached for the first time
            if i > TargetTemp:
                FirstStepPassed = True

        else:
            # Use PID threshold,
            # not temperature threshold

            # Compare current PID output
            # to PID threshold
            if output <= PID_thres:

                # Turn OFF the relay and the heater
                GPIO.output(18,GPIO.LOW)

                # Print current temperature and
                # heater status
                print(i, "Degree celcius, ", "Heater OFF, ", "PID: ", output)

                # Update current temperature display
                CurrTempDisplay.value = str(round(i,2))
                app.update()

            else:
                # Turn ON the relay and the heater
                GPIO.output(18,GPIO.HIGH)

                # Print current temperature and
                # heater status
                print(i, "Degree celcius, ", "Heater OFF, ", "PID: ", output)

                # Update current temperature display
                CurrTempDisplay.value = str(round(i,2))
                app.update()

    else:
        # Only update current temperature

        # Read temperature
        i = read_temp()

        # Update current temperature display
        CurrTempDisplay.value = str(round(i,2))

##### Main Function (End) #####

##### Initialize temperature sensor #####

os.system('modprobe w1-gpio')
os.system('modprobe w1-therm')

# These are default directories for
# the temperature sensor
# (DON'T CHANGE)
base_dir = '/sys/bus/w1/devices/'
device_folder = glob.glob(base_dir + '28*')[0]
device_file = device_folder + '/w1_slave'

##### Initialize Relay GPIO pin (Pin 18) #####

GPIO.setmode(GPIO.BCM)
GPIO.setwarnings(False)
GPIO.setup(18,GPIO.OUT)

##### Initialize global variables #####

# For time logging
StartSeconds = time.time()
CurrSeconds = time.time()

# For logging and saving data
now = datetime.now()
filename = str(now)
filename = filename.replace(" ", "_")
filename = filename.replace(":", ".")
curr_time = str(now)
curr_time = curr_time.replace(" ", "_")
curr_time = curr_time.replace(":", ".")
data = [[curr_time, 0, 0, 0]]
df = pd.DataFrame(data, columns = ['DateTime', 'Time (s)', 'temp', 'TargetTemp'])

# PID variables
P = 1
I = 0
D = 0

PID_thres = 0.25

pid = PID.PID(P, I, D)

# Other variables
TargetTemp = 0
screen_capture = ""
SystemOn = False

# For checking if target temp has been
# reached for the first time or not. In this case,
# do not use PID tolerance threshold.
# Use a temperature tolerance threshold based
# on the target temperature. After it has been
# reached for the first time,
# use PID tolerance threshold
FirstStepPassed = False

###### Configure app #####

app = App("Heater system", height=400, width=500)

##### Display Boxes #####

TargetTempDisplayBox = Box(app, border=True, height=50, width=350)
TargetTempDisplayTitle = Text(TargetTempDisplayBox, text="Target Temperature")
TargetTempDisplay = Text(TargetTempDisplayBox, text="")

CurrTempDisplayBox = Box(app, border=True, height=50, width=350)
CurrTempDisplayTitle = Text(CurrTempDisplayBox, text="Current Temperature")
CurrTempDisplay = Text(CurrTempDisplayBox, text="")

StatusDisplayBox = Box(app, border=True, height=50, width=350)
StatusDisplay = Text(StatusDisplayBox, text="Standby")

##### Numbers Buttons #####

padding = 10
text_size = 10
height = 5
width = 5

numbersBox = Box(app, layout="grid", align="top", height=230, width=350)

one = PushButton(numbersBox, text="1", command=lambda:screenCapture(1) , grid=[0,0], height = height, width = width, padx = padding, pady = padding)
two = PushButton(numbersBox, text="2", command=lambda:screenCapture(2) , grid=[1,0], height = height, width = width, padx = padding, pady = padding)
three = PushButton(numbersBox, text="3", command=lambda:screenCapture(3) , grid=[2,0], height = height, width = width, padx = padding, pady = padding)
four = PushButton(numbersBox, text="4", command=lambda:screenCapture(4) , grid=[3,0], height = height, width = width, padx = padding, pady = padding)
five = PushButton(numbersBox, text="5", command=lambda:screenCapture(5) , grid=[4,0], height = height, width = width, padx = padding, pady = padding)
six = PushButton(numbersBox, text="6", command=lambda:screenCapture(6) , grid=[0,1], height = height, width = width, padx = padding, pady = padding)
seven = PushButton(numbersBox, text="7", command=lambda:screenCapture(7) , grid=[1,1], height = height, width = width, padx = padding, pady = padding)
eight = PushButton(numbersBox, text="8", command=lambda:screenCapture(8) , grid=[2,1], height = height, width = width, padx = padding, pady = padding)
nine = PushButton(numbersBox, text="9", command=lambda:screenCapture(9) , grid=[3,1], height = height, width = width, padx = padding, pady = padding)
zero = PushButton(numbersBox, text="0", command=lambda:screenCapture(0) , grid=[4,1], height = height, width = width, padx = padding, pady = padding)
one.text_size = text_size
two.text_size = text_size
three.text_size = text_size
four.text_size = text_size
five.text_size = text_size
six.text_size = text_size
seven.text_size = text_size
eight.text_size = text_size
nine.text_size = text_size
zero.text_size = text_size

##### Input/Output Buttons #####

ioBox = Box(app, layout="grid", align="top", height=200,width=350)

start_button = PushButton(ioBox, text="START", command=StartSystem, grid=[0,0], width = 5, height = 5)
start_button.bg = (0,200,0)

stop_button = PushButton(ioBox, text="STOP", command=StopSystem, grid=[1,0], width = 5, height = 5)
stop_button.bg = "yellow"

delete_button = PushButton(ioBox, text="DEL", command=backSpace, grid=[2,0], width = 5, height = 5)

exit_button = PushButton(ioBox, text="EXIT", command=exitApp, grid=[4,0], width = 5, height = 5)

shutdown_button = PushButton(ioBox, text="SHUT\nDOWN", command=Shutdown, grid=[5,0], width = 5, height = 5)
shutdown_button.bg = "red"

##### Run main code every 1 second #####

display.repeat(1000, mainRun)
app.tk.attributes("-fullscreen",True)
app.display()
```

```
# PID.py

# ==============================================================================
# This file is part of IvPID.
# Copyright (C) 2015 Ivmech Mechatronics Ltd. <>
#
# IvPID is free software: you can redistribute it and/or modify
# it under the terms of the GNU General Public License as published by
# the Free Software Foundation, either version 3 of the License, or
# (at your option) any later version.
#
# IvPID is distributed in the hope that it will be useful,
# but WITHOUT ANY WARRANTY; without even the implied warranty of
# MERCHANTABILITY or FITNESS FOR A PARTICULAR PURPOSE.  See the
# GNU General Public License for more details.
#
# You should have received a copy of the GNU General Public License
# along with this program.  If not, see <http://www.gnu.org/licenses/>.

# title           :PID.py
# description     :python pid controller
# author          :Caner Durmusoglu
# date            :20151218
# version         :0.1
# notes           :
# python_version  :2.7
# ==============================================================================

"""Ivmech PID Controller is simple implementation of a Proportional-Integral-Derivative (PID) Controller in the Python Programming Language.
More information about PID Controller: http://en.wikipedia.org/wiki/PID_controller
"""
import time

class PID:
    """PID Controller
    """

    def __init__(self, P=0.2, I=0.0, D=0.0, current_time=None):

        self.Kp = P
        self.Ki = I
        self.Kd = D

        self.sample_time = 0.00
        self.current_time = current_time if current_time is not None else time.time()
        self.last_time = self.current_time

        self.clear()

    def clear(self):
        """Clears PID computations and coefficients"""
        self.SetPoint = 0.0

        self.PTerm = 0.0
        self.ITerm = 0.0
        self.DTerm = 0.0
        self.last_error = 0.0

        # Windup Guard
        self.int_error = 0.0
        self.windup_guard = 20.0

        self.output = 0.0

    def update(self, feedback_value, current_time=None):
        """Calculates PID value for given reference feedback

        .. math::
            u(t) = K_p e(t) + K_i \int_{0}^{t} e(t)dt + K_d {de}/{dt}

        .. figure:: images/pid_1.png
           :align:   center

           Test PID with Kp=1.2, Ki=1, Kd=0.001 (test_pid.py)

        """
        error = self.SetPoint - feedback_value

        self.current_time = current_time if current_time is not None else time.time()
        delta_time = self.current_time - self.last_time
        delta_error = error - self.last_error

        if (delta_time >= self.sample_time):
            self.PTerm = self.Kp * error
            self.ITerm += error * delta_time

            if (self.ITerm < -self.windup_guard):
                self.ITerm = -self.windup_guard
            elif (self.ITerm > self.windup_guard):
                self.ITerm = self.windup_guard

            self.DTerm = 0.0
            if delta_time > 0:
                self.DTerm = delta_error / delta_time

            # Remember last time and last error for next calculation
            self.last_time = self.current_time
            self.last_error = error

            self.output = self.PTerm + (self.Ki * self.ITerm) + (self.Kd * self.DTerm)

    def setKp(self, proportional_gain):
        """Determines how aggressively the PID reacts to the current error with setting Proportional Gain"""
        self.Kp = proportional_gain

    def setKi(self, integral_gain):
        """Determines how aggressively the PID reacts to the current error with setting Integral Gain"""
        self.Ki = integral_gain

    def setKd(self, derivative_gain):
        """Determines how aggressively the PID reacts to the current error with setting Derivative Gain"""
        self.Kd = derivative_gain

    def setWindup(self, windup):
        """Integral windup, also known as integrator windup or reset windup,
        refers to the situation in a PID feedback controller where
        a large change in setpoint occurs (say a positive change)
        and the integral terms accumulates a significant error
        during the rise (windup), thus overshooting and continuing
        to increase as this accumulated error is unwound
        (offset by errors in the other direction).
        The specific problem is the excess overshooting.
        """
        self.windup_guard = windup

    def setSampleTime(self, sample_time):
        """PID that should be updated at a regular interval.
        Based on a pre-determined sampe time, the PID decides if it should compute or return immediately.
        """
        self.sample_time = sample_time
```
